# ATP8B1–TMEM30B Flippase Activity Maintains Stereocilia Lipid Asymmetry Required for Hearing

**DOI:** 10.64898/2026.02.12.705475

**Authors:** Henry N. De Hoyos, Sihan Li, Jun-Sub Im, Alyssa Luz-Ricca, Takesha Foster, Betsy Szeto, Rachel Jonas, Emma Kim, Nikhil Amin, Jung-Bum Shin

## Abstract

Sensory hair cells convert sound-induced vibrations into electrical signals through a process called mechano-electrical transduction (MET). While the protein components of the MET complex are well studied, increasing evidence indicates that MET channel properties are significantly modulated by the surrounding lipid bilayer. The asymmetric distribution of membrane lipids between the inner and outer membrane leaflets is well established to shape membrane mechanics. The recent discovery that the core MET components TMC1 and TMC2 also act as lipid scramblases suggests a direct role for membrane lipid asymmetry in the dynamic shaping of auditory transduction. Because scramblase activity of TMC1/2 disrupts lipid asymmetry, we hypothesized that an opposing flippase may be required to restore and maintain lipid asymmetry. Here, we identify the P4-ATPase ATP8B1 and its chaperone TMEM30B as selectively expressed in outer hair cells (OHCs), enriched in stereocilia, and upregulated following the onset of MET and hearing. Loss of either protein results in elevated auditory brainstem response (ABR) thresholds, phosphatidylserine (PS) externalization, and rapid hair-cell degeneration, demonstrating that lipid homeostasis is crucial for OHC survival. Together, these findings establish ATP8B1 and TMEM30B as key regulators of membrane lipid asymmetry in sensory hair cells and establish TMEM30B as a novel deafness gene.

**Impact Statement:** The ATP8B1–TMEM30B flippase complex maintains outer hair-cell membrane asymmetry, which is essential for hair cell survival and auditory function.

## Introduction

Hearing depends on the ability of mechanosensory hair cells to convert sound-evoked mechanical forces into electrical signals (Hudspeth 1989; Gillespie and Müller 2009; Fettiplace and Kim 2014). Sound vibrations transmitted through the middle ear displace the basilar membrane and deflect the stereocilia bundles that crown auditory hair cells. This deflection modulates tension on tip links which are filamentous connectors that couple adjacent stereocilia and directly gates mechanotransduction (MET) channels. The architecture of the hair bundle, including the graded heights of stereocilia rows and the polarity of the MET machinery, is essential for tuning the sensitivity and dynamic range of auditory signaling. Many of the protein components of the MET complex have now been identified. At the core of this machinery is a force-transmission complex that couples the extracellular tip link to the MET channel complex. Tip links are formed by cadherin 23 and protocadherin 15 (Pickles et al. 1984; Assad et al. 1991; Siemens et al. 2004; Kazmierczak et al. 2007; Ahmed et al. 2006), which relay mechanical force to the MET channel through a network of associated proteins, including LHFPL5 (Xiong et al. 2012), CIB2 (Riazuddin et al. 2012; Giese et al. 2017), and TMIE (Zhao et al. 2014). TMC1 and its paralog TMC2 are proposed to constitute the ion-conducting components of the MET channel, with TMC1 serving as the dominant pore-forming subunit in mature mammalian hair cells (Kawashima et al. 2011; Pan et al. 2013, 2018; Assad et al. 1991; Pickles et al. 1984). Intriguingly, TMC1 and TMC2 share structural similarity with TMEM16 family lipid scramblases, of which TMEM16A and B function as ion channels (Le and Yang 2021). More broadly, TMEM16, TMC, and TMEM63/OSCA are members of a structurally related superfamily of membrane proteins that includes ion channels and lipid scramblases (Medrano-Soto et al. 2018). Consistent with this relationship, TMC1 and TMC2 were shown to possess lipid-scrambling activity in addition to their proposed ion-conducting function (Ballesteros et al. 2018; Ballesteros and Swartz 2022; Clark et al. 2024; Peineau et al. 2025; George and Ricci 2025). Cryo-EM structures of *C. elegans* homologues of TMC1 reveal that TMC forms a dimer associated with TMIE and calcium-binding accessory proteins and engages extensively with the surrounding lipid bilayer (Jeong et al. 2022; Clark et al. 2024). These structures resolve conserved lipid-like densities at defined positions around the complex, including a lipid-filled cavity at the TMIE-TMC interface and sterol-like densities near the dimer interface (Jeong et al. 2022; Clark et al. 2024). Such lipid-mediated interactions appear to stabilize subunit assembly and may influence conformational changes during channel gating. Together, these findings suggest that the membrane is an active component of the MET complex, with lipid-protein interactions poised to shape MET properties including sensitivity, resting open probability and adaptation.

These observations further imply that MET function may require a specialized lipid environment within stereocilia, whose composition and organization are distinct from the surrounding apical membrane. However, the molecular mechanisms, including the lipid-modifying enzymes, transport pathways, and regulatory factors, that establish, maintain, and dynamically regulate this unique lipid milieu are unknown.

Lipids are not uniformly distributed across the plasma membrane (Bretscher 1972). Instead, eukaryotic membranes maintain a highly asymmetric distribution of phospholipids between their inner and outer leaflets (Cooper 2000; Lorent et al. 2020). Phosphatidylserine (PS) and phosphatidylethanolamine (PE) are normally enriched on the cytoplasmic leaflet, whereas phosphatidylcholine and sphingomyelin predominate externally (Cooper 2000; Lorent et al. 2020). This asymmetry is actively maintained by ATP-dependent flippases and floppases and disrupted by scramblases (Nagata et al. 2020).

Possibly best studied is the asymmetric distribution of PS. Loss of PS asymmetry is widely recognized as a potent cellular signal, as PS is normally confined to the cytoplasmic leaflet by ATP-dependent flippases (Nagata et al. 2020). Its externalization is best known as a hallmark of apoptosis, where it functions as an “eat-me” signal that promotes rapid clearance of dying cells (Nagata et al. 2010; Suzuki et al. 2010). However, PS exposure is not exclusively associated with cell death. In immune cells and platelets, transient PS externalization accompanies physiological activation states and serves signaling or catalytic roles, illustrating that regulated loss of asymmetry can be functionally repurposed (Shin and Takatsu 2020). PS exposure has also been implicated in cell-cell fusion events, where localized scrambling appears to facilitate membrane remodeling and lower energetic barriers to cell-cell fusion (Whitlock and Chernomordik 2021).

Beyond signaling, lipid asymmetry has significant biophysical consequences for the membrane. Asymmetric enrichment of charged, unsaturated lipids such as PS and PE in the inner leaflet, opposed by phosphatidylcholine, sphingomyelin- and cholesterol-rich outer leaflets, creates differences in lipid packing, order, and viscosity across the bilayer (Reddy et al. 2012; Hossein and Deserno 2020; Doktorova et al. 2020; Kakuda et al. 2022; X. Li et al. 2024; Wang et al. 2025). This asymmetry strongly influences membrane tension, curvature stress, and bending rigidity, parameters that govern how membranes deform and how embedded proteins respond to force (Lorent et al. 2020; Hossein and Deserno 2020; Wang et al. 2025).

In this context, the scramblase activity associated with TMC proteins raises an important question: does TMC-mediated lipid scrambling represent an unintended byproduct of MET, or does it serve a specific functional role? Initial models proposed that scrambling might facilitate adaptive membrane remodeling or ectosome shedding (Ballesteros and Swartz 2022). Given the exquisite sensitivity of MET channel function to the surrounding lipid environment, it is also hypothesized that regulated scrambling directly modulates MET gating by altering membrane mechanical properties (George and Ricci 2025). Regardless of its precise physiological benefit, sustained lipid scrambling would be incompatible with cellular homeostasis, as aberrant PS exposure and loss of membrane asymmetry are broadly harmful for the cell (Ballesteros and Swartz 2022; Beurg et al. 2025). Thus, mechanisms that actively restore lipid asymmetry are likely essential in hair cells.

P4-ATPases, the canonical phospholipid flippases, are prime candidates for maintaining this asymmetry (Segawa et al. 2016). These enzymes selectively transport PS and other phospholipids from the outer to the inner leaflet and require CDC50 family accessory subunits (also known as TMEM30A, TMEM30B, and TMEM30C) for proper folding, trafficking, and activity (Lenoir et al. 2009; Bryde et al. 2010; van der Velden et al. 2010; Segawa et al. 2018) (**Fig.1A**). While TMEM30A is ubiquitously expressed and supports multiple P4-ATPases, TMEM30B exhibits more restricted tissue distribution and remains poorly characterized (Segawa et al. 2018; T. Li et al. 2021; Xing et al. 2023). TMEM30C is known to only be expressed in mammalian testis (Osada et al. 2007). Transcriptomic analyses indicate that TMEM30B is expressed in the inner ear (Liu et al. 2018; Scheffer et al. 2015; Kolla et al. 2020; Orvis et al. 2021), and a recent hearing loss screen in a cohort of 3006 mouse knockout (KO) strains reported that *Tmem30b* KO mice develop hearing loss (Bowl et al. 2017). However, the cellular and molecular basis of the mechanism underlying the hearing loss, and the developmental regulation and subcellular localization of TMEM30B in hair cells remained unknown.

**Figure 1:**
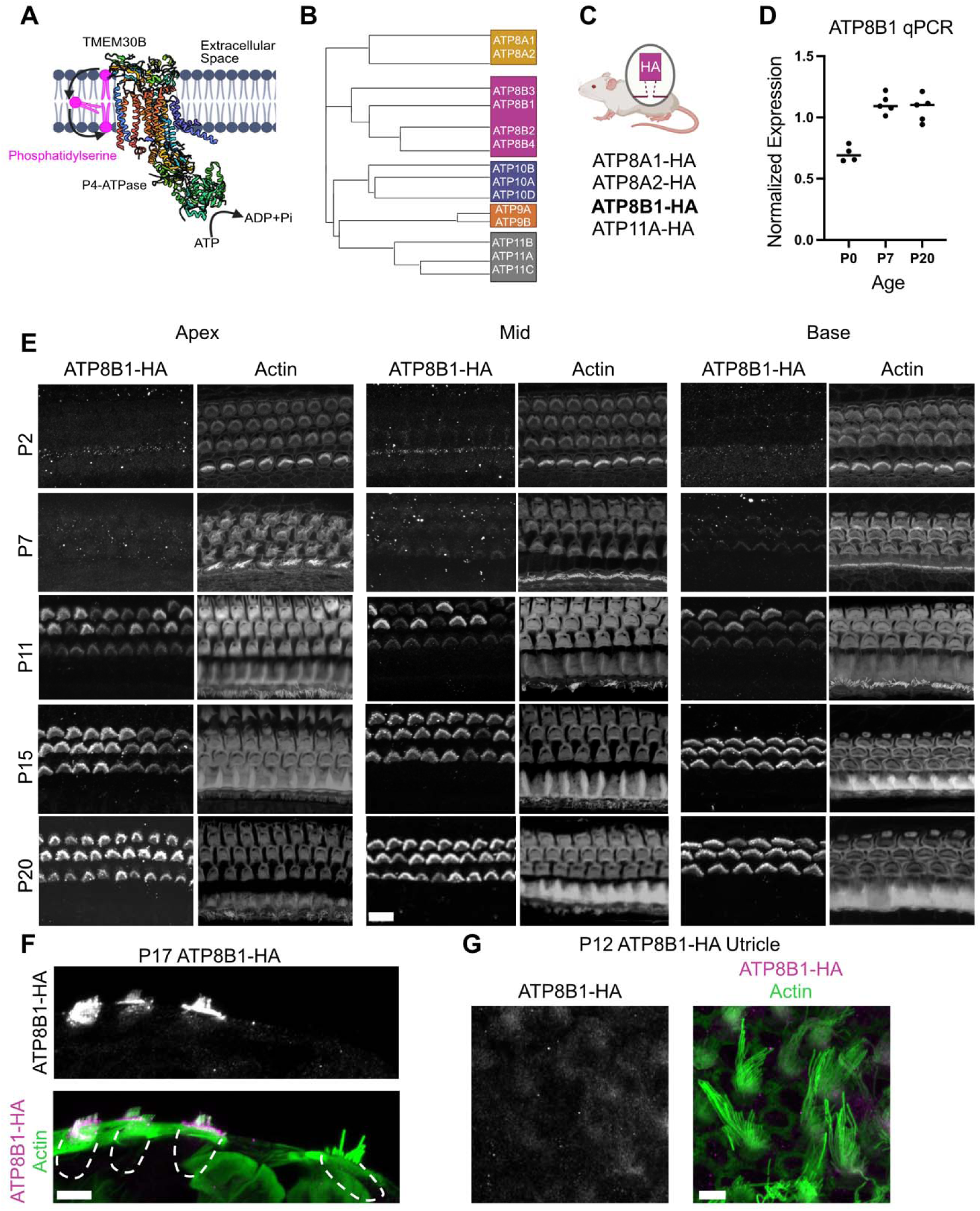
ATP8B1 is a P4-ATPase flippase selectively expressed in outer hair cells. **(A)** Schematic illustrating ATP8B1 and its obligate chaperone TMEM30B embedded in the plasma membrane (adopted from PDB, 8OX4) (Dieudonné et al. 2023). Created in BioRender. Shin, J. (2026) https://BioRender.com/vo62i9e. (**B)** Cladogram of the 14 P4-ATPase family members expressed in mice. Created in BioRender. Shin, J. (2026) https://BioRender.com/owpxh5c. **(C)** Of the 14 P4-ATPases, four were detected in hair cells, and HA knock-in mouse lines were generated for each. ATP8B1-HA (bold) was the only flippase found to be expressed in outer hair cells Created in BioRender. Shin, J. (2026) https://BioRender.com/ed4b3cb. **(D)** qPCR analysis of *Atp8b1* expression at P0, P7, and P20. Four to five cochleae were analyzed per age group. **(E)** ATP8B1-HA Immunofluorescence across development and along the cochlear tonotopic axis in the organ of Corti. Organ of Corti were stained for anti-HA and phalloidin for F-actin. **(F)** Orthogonal view of ATP8B1-HA immunofluorescence in a P17 organ of Corti. A Compresstome section was used to facilitate imaging of the hair bundle plane. Scale bars: 5 μm **(G)** ATP8B1-HA immunofluorescence in P12 utricle showing minimal expression. Scale bars 10 μm.

Here, we investigate the roles of the flippase ATP8B1 and its accessory subunit TMEM30B in auditory hair cells. We show that both proteins are highly enriched in stereocilia and the apical membrane. Using genetic KO mouse models, we demonstrate that loss of ATP8B1 leads to loss of lipid membrane asymmetry, early-onset hair cell pathology and rapid hearing decline. *Tmem30b* KO mice phenocopy these defects. Together, our findings identify ATP8B1 and TMEM30B as essential regulators of membrane lipid asymmetry in OHCs and reveal a previously underappreciated role for lipid flippase pathways in hair cell function and survival.

## Results

### ATP8B1 is specifically targeted to outer hair cell stereocilia

Of the 14 P-type ATPase flippases conserved between mice and humans, nine localize to the plasma membrane (Shin and Takatsu 2025), and transcriptomic analyses identify four as being expressed in hair cells (Mark et al. 2013; Liu et al. 2018; Kolla et al. 2020). Recent work has implicated several phospholipid flippases, including ATP8B1, ATP8A2, and ATP11A in auditory dysfunction (Coleman et al. 2014; Pater et al. 2022; Stapelbroek et al. 2009). Because available antibodies lacked sufficient specificity, we generated HA knock-in (KI) mouse lines for ATP8A1, ATP8A2, ATP8B1 and ATP11A (**Fig. 1B-C**). Importantly, ATP8B1-HA mice exhibited normal hearing thresholds compared to C57BL/6J controls at P30, indicating that insertion of the HA tag did not impair auditory function (**Figure 1-Sup-1**). Together, these data support the use of the ATP8B1-HA knock-in allele for localization studies. Among these candidates, ATP8B1 was the only flippase specifically enriched in the hair bundle of OHCs, consistent with previous studies reporting ATP8B1-dependent hair-cell degeneration and hearing loss (Stapelbroek et al. 2009). Remarkably, no ATP8B1-HA immunoreactivity was detected in IHCs or their hair bundles. We performed a developmental time course of expression. Organs of Corti from ATP8B1-HA mice were imaged at key stages spanning hair-cell maturation: P2 (around the onset of MET activity), P7 (MET established but prior to hearing onset), P12 (near hearing onset), P17 and P20 (mature auditory epithelium) (**Fig. 1E**). Across this timeline, ATP8B1 became progressively enriched in OHC stereocilia, coinciding with maturation of MET function and hearing onset. Protein expression patterns matched transcript levels, as qPCR analysis for *Atp8b1* demonstrated a developmental trajectory consistent with our immunofluorescence data (**Fig. 1D**).

**Figure 1-Supplement-1:**
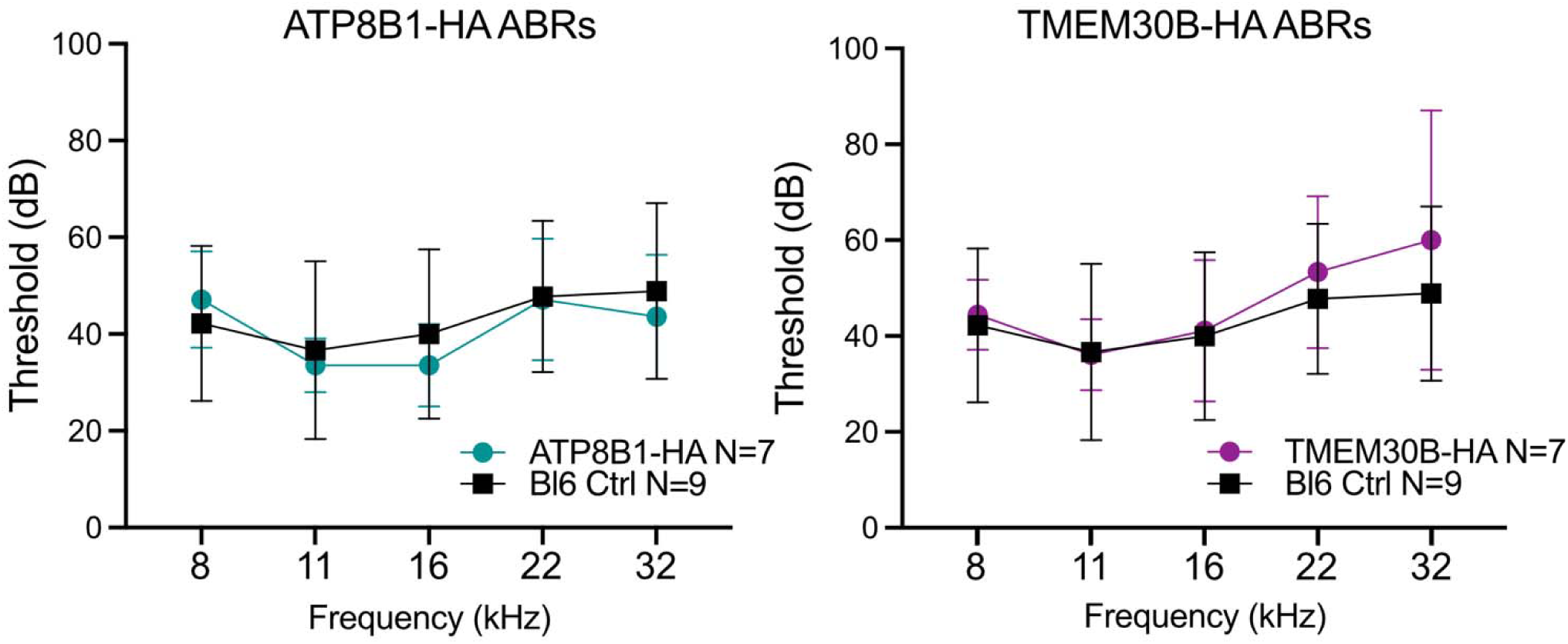
ATP8B1-HA and TMEM30B-HA mice have normal hearing. Auditory brainstem response (ABR) thresholds at P30 in at ATP8B1-HA and TMEM30B-HA (ATP8B1-HA, N = 7; TMEM30B-HA, N =7, Bl6 WT ctrl, N=9). ABR thresholds were not significantly elevated at all tested frequencies (8, 11, 16, 22, and 32 kHz; Mann–Whitney U test, NS)

Finally, we assessed whether ATP8B1 is restricted to the stereocilia or distributed throughout the hair cell. ATP8B1-HA localized almost exclusively to OHC bundles and the apical membrane, with minimal staining in the remaining plasma membrane (**Fig. 1F**). In contrast, staining of the utricle revealed only weak ATP8B1-HA signal in vestibular hair cells, markedly lower than that observed in OHCs (**Fig. 1G**). Together, these data establish ATP8B1 as a tightly localized flippase that becomes strongly bundle-enriched during OHC maturation, with substantially lower expression in vestibular hair cells, and showed no detectable signal in IHCs.

We next investigated the precise subcellular localization of ATP8B1 in OHC stereocilia. OHCs oriented in profile were identified and imaged. Quantitative fluorescence intensity profiling of ATP8B1-HA and actin along both short (row 2) and tall (row 1) stereocilia revealed that ATP8B1 is present in all stereocilia rows. Notably, ATP8B1 is not uniformly distributed along the length of the stereocilia; instead, it is enriched at the stereociliary base, with signal intensity progressively decreasing toward the tips (**Fig. 2A, B, C, D**).

**Figure 2:**
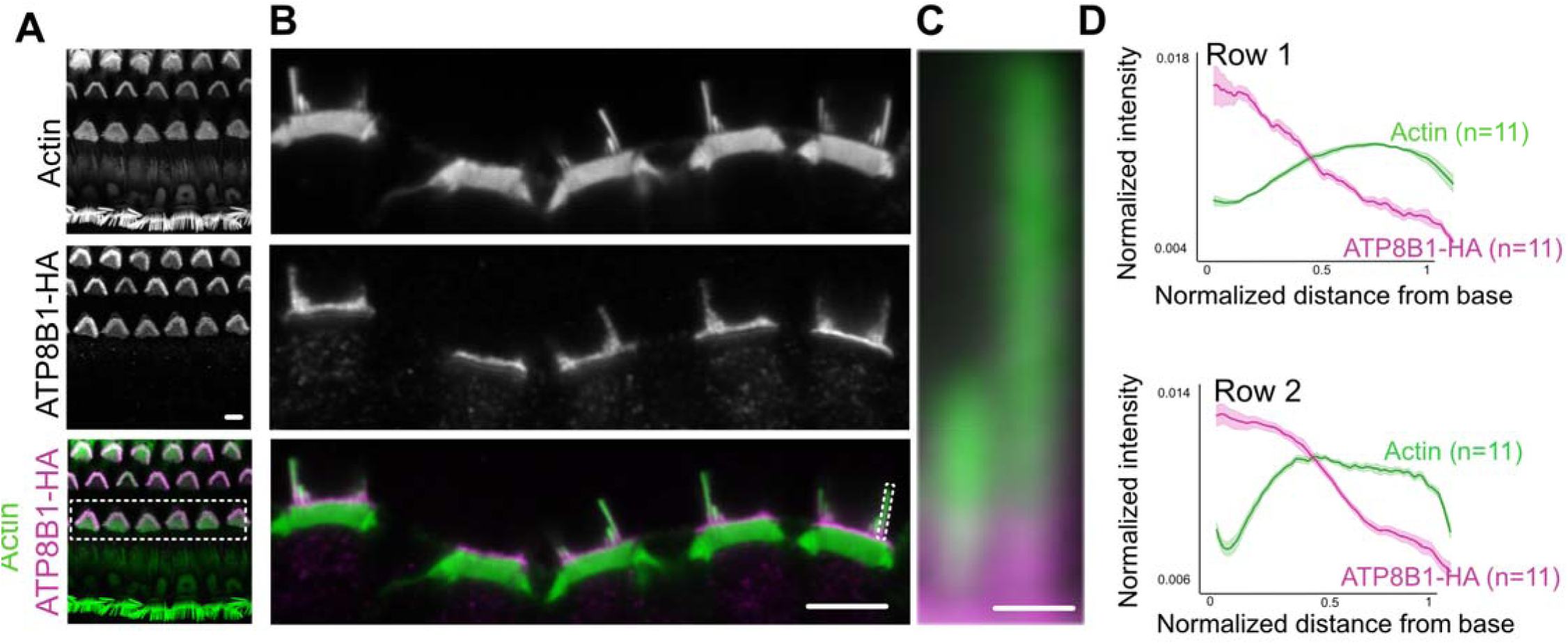
ATP8B1 is enriched toward the base of stereocilia. **(A)** ATP8B1-HA immunofluorescence showing OHC specific expression. (**B,C)** ATP8B1-HA IF is enriched towards the stereocilia base and the apical hair cell membrane. **(D)** Average normalized fluorescence intensity profiles of actin and ATP8B1 along short (row 2) and long (row 1) stereocilia. Position is normalized from 0 (base) to 1 (tip), and intensities are normalized to total signal per stereocilium. Shaded regions indicate SEM. (Scale bar: 5 μm and 0.5 μm in stereocilia inset)

### Loss of ATP8B1 leads to early onset hearing loss and rapid OHC degeneration

To define the role of ATP8B1 in auditory function, we generated a *Atp8b1* knockout (KO) mouse (19 bp deletion allele disrupting the translational start). Examination of the cochlea revealed a rapid and progressive loss of OHCs, beginning in the basal turn by postnatal day 17 and extending apically over the next several days (**Fig. 3A, B**). By P20, only a small number of basal OHCs remained (**Fig. 3B, C**). In contrast, IHCs, in which we did not detect ATP8B1, were preserved (**Fig. 3D**), indicating a selective requirement for ATP8B1 in OHC survival. Consistent with this cellular phenotype, *Atp8b1* KO mice exhibited early-onset hearing loss. Auditory brainstem response (ABR) thresholds and distortion product otoacoustic emissions (DPOAEs) were significantly elevated across all tested frequencies at P20 compared with littermate controls, consistent with loss of OHC function (**Fig. 3E**). This phenotype mirrors that previously reported for an independent *Atp8b1* point-mutation allele (Stapelbroek et al. 2009). Together, these findings demonstrate that ATP8B1 is essential for OHC survival and auditory function.

**Figure 3.**
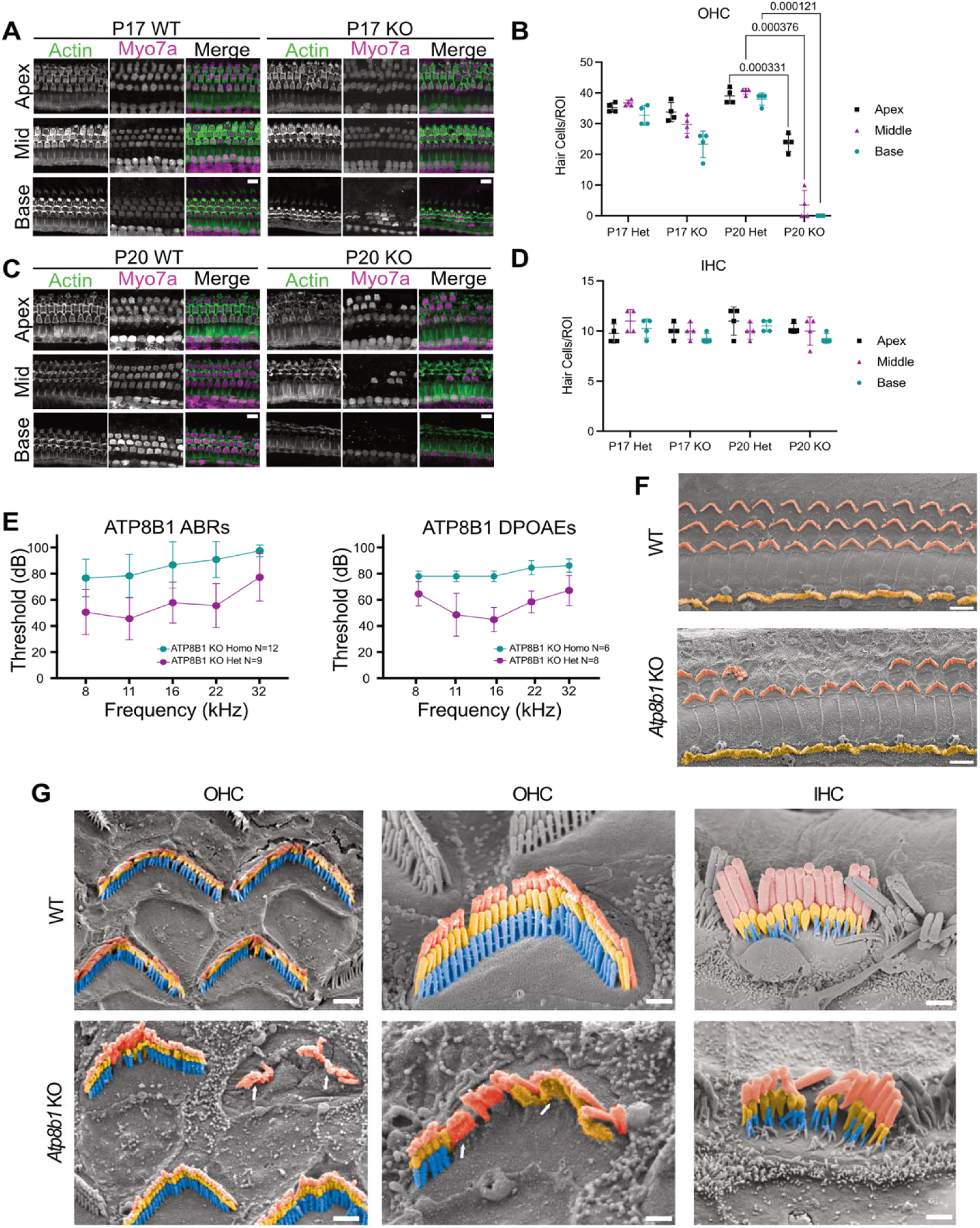
Loss of ATP8B1 causes progressive OHC bundle degeneration, hair cell loss, and elevated auditory thresholds. **(A,B)** Representative immunofluorescence images of WT and *Atp8b1* KO cochleae at P17 and P20 stained for MYO7A (hair cells) and phalloidin (F-actin). OHC loss is first observed in the basal turn of *Atp8b1* KO cochleae. Scale bar, 10 μm. **(C)** Quantification of OHC numbers in WT and *Atp8b1* KO cochleae at P17 and P20 across apical, middle, and basal turns (n = 3–5 cochleae per group). At P20, OHC loss was significant in all tonotopic regions in KO mice compared with WT (Welch’s t-test, *p* < 0.001), whereas no significant differences were observed at P17. **(D)** Quantification of IHC survival in WT and *Atp8b1* KO cochleae at P17 and P20 across apical, middle, and basal turns (n = 3–5 cochleae per group). No significant differences were detected between genotypes at either age (Welch’s t-test). **(E)** Auditory brainstem response (ABR) thresholds at P30 are significantly elevated in *Atp8b1* KO mice compared with littermate controls (KO, N = 12; WT, N = 9). Distortion product otoacoustic emissions (DPOAE) thresholds at P20 are also significantly elevated, (KO, N = 6; WT, N = 8). ABR thresholds were increased at all tested frequencies (8, 11, 16, 22, and 32 kHz; Mann–Whitney U test, *p* < 0.001). DPOAE thresholds were increased at all tested frequencies. (8, 11, 16, 22, and 32 kHz; Mann–Whitney U test, (*p<.01,* 8khz), *p* < 0.001, 11-32khz) **(F)** Scanning electron microscopy (SEM) images of WT and *Atp8b1* KO organs of Corti at P20. OHCs are pseudo-colored orange and IHCs yellow. Scale bars: 5 μm. **(G)** High-magnification SEM images at P20 reveal stereocilia fusion and OHC loss in *Atp8b1* KO mice, features absent in WT. Arrows point out areas of stereocilia fusion and loss. IHCs remain morphologically intact. Scale bars: 1 μm (left), 0.5 μm (middle and right).

Following the observed hair cell loss of *Atp8b1* KO mice, we performed scanning electron microscopy (SEM) to assess hair-bundle morphology and cell integrity (**Fig. 3F**). *Atp8b1* KO OHCs showed marked morphological deterioration, characterized by reduced transducing stereocilia number and occasional fusion along the edges of stereocilia rows, a known pathological feature associated with progressive hearing loss (**Fig. 3G**). In contrast, IHCs did not exhibit detectable bundle abnormalities or cell loss, consistent with our immunohistochemistry results. Notably, stereocilia degeneration and fusion in the KO mice occurred prior to overt hair cell loss, indicating that disruption of bundle architecture is an early consequence of ATP8B1 deficiency.

### TMEM30B expression and localization mirrors that of ATP8B1

The targeting and function of P-type ATPase flippases requires association with β-subunits TMEM30A or TMEM30B. To determine which partners operate with ATP8B1 in OHCs, we created HA-tagged KI mice for both proteins. TMEM30B, but not TMEM30A (data not shown), was expressed in hair cells. TMEM30B-HA mice exhibited normal hearing thresholds compared with C57BL/6J controls at P30, indicating that insertion of the HA tag did not disrupt auditory function (**Figure 1-Sup-1**). Organs of Corti from TMEM30B-HA mice were imaged at P2, P7, P12, P17 and P20 (**Fig. 4A**). Across this series, TMEM30B-HA showed a progressive enrichment in stereocilia, paralleling the maturation of the MET apparatus and the onset of hearing, similar to ATP8B1. We next studied the subcellular distribution of TMEM30B. While ATP8B1-HA was almost exclusively confined to OHC bundles (**Fig. 1 and 2**), TMEM30B-HA was enriched in the stereocilia compartment, but also weakly in the cell body (**Fig. 4B**). Like ATP8B1-HA, TMEM30B-HA showed modest labeling of utricular hair bundles at P17 (**Fig. 4C**) and was excluded from IHC bundles. Together, these findings identify ATP8B1 and TMEM30B as a tightly co-localized flippase complex that becomes progressively OHC bundle-specific during auditory maturation.

**Figure 4.**
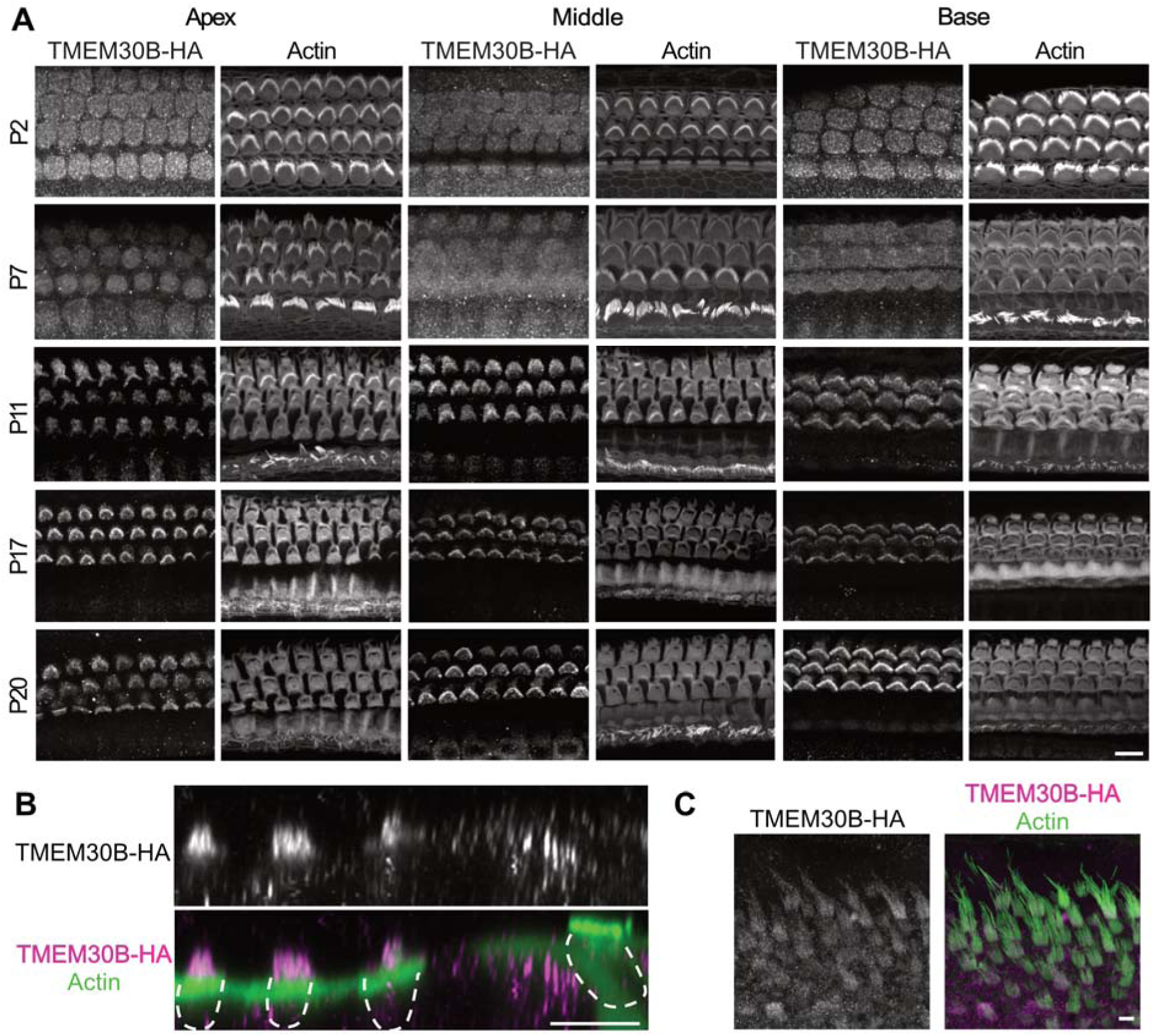
The P4-ATPase chaperone TMEM30B is enriched in OHC bundles. **(A)** Immunofluorescence images of TMEM30B-HA across development and along the cochlear tonotopic axis in the organ of Corti. Organ of Corti were stained for anti-HA and phalloidin for F-actin. Scale bars 10 μm. **(B)** Orthogonal view of TMEM30B-HA immunofluorescence of a P17 organ of Corti. Scale bars 10 μm. **(C)** Immunofluorescence images of a P17 TMEM30B-HA utricle showing moderate expression in utricular hair cells. Scale bars 10 μm.

### TMEM30B is required for hearing and OHC maintenance

Next, we generated *Tmem30b* KO mice harboring a 54 bp deletion disrupting the translational start. Similar to *Atp8b1* KO mice, *Tmem30b* KO mice exhibited rapid OHC degeneration, beginning in the basal turn at P17 and advancing apically over the following days. By P20, only a small number of basal OHCs remained (**Fig. 5A, B**). In contrast, IHCs were preserved, closely mirroring the *Atp8b1* KO phenotype and indicating that loss of TMEM30B selectively compromises OHC survival (**Fig. 5C, D**). Consistent with this cellular phenotype, *Tmem30b* KO mice exhibited early-onset hearing loss. Auditory brainstem response (ABR) thresholds and distortion product otoacoustic emissions (DPOAEs) were significantly elevated across all tested frequencies at P20 compared with littermate controls, consistent with loss of OHC function (**Fig. 5E**). Together, these findings confirm TMEM30B as a deafness gene, in agreement with an auditory phenotype previously reported by the IMPC (International Mouse Phenotyping Consortium) (Bowl et al. 2017).

**Figure 5.**
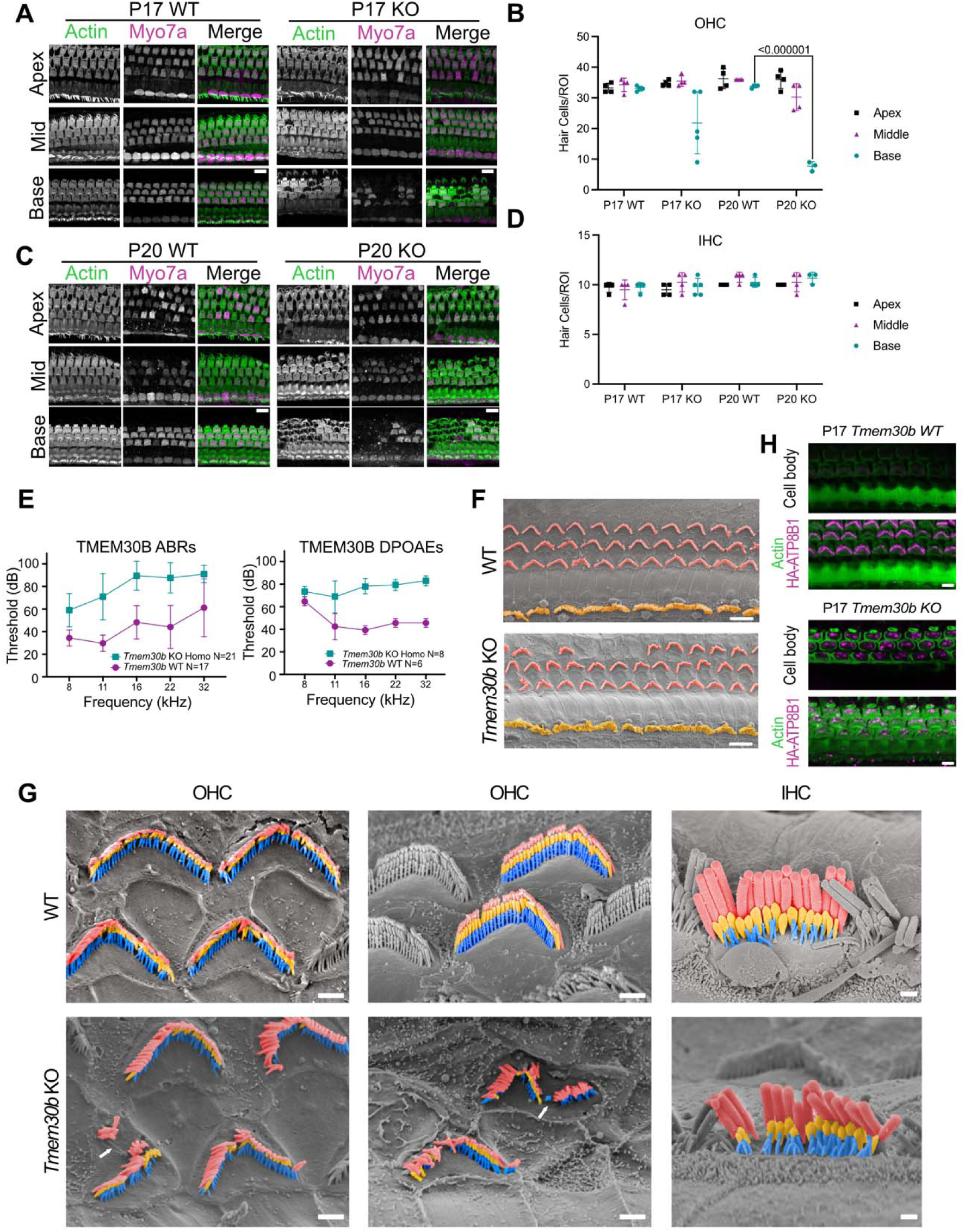
Loss of TMEM30B causes progressive OHC degeneration, hair cell loss, and elevated auditory thresholds. **(A,B)** Representative Immunofluorescence images of WT and *Tmem30b* KO cochleae at P17 and P20, stained for MYO7A (hair cells) and phalloidin (F-actin). OHC loss is first detected in the basal turn of *Tmem30b* KO cochleae at P17, and pronounced OHC loss in the basal turn of KO cochleae at P20.. Scale bar, 10 μm. **(C)** Quantification of OHC numbers in WT and *Tmem30b* KO cochleae at P17 and P20 across apical, middle, and basal turns (n = 3–5 cochleae per group). At P20, OHC loss was significant in the basal region of KO mice compared with WT (Welch’s *t*-test, *p* < 0.001), whereas no significant differences were observed at P17. **(D)** Quantification of IHC numbers in WT and *Tmem30b* KO cochleae at P17 and P20 across apical, middle, and basal turns (n = 3–5 cochleae per group). No significant differences were detected between genotypes at either age (Welch’s *t*-test). **(E)** Auditory brainstem response (ABR) thresholds at P30 are significantly elevated in *Tmem30b* KO mice compared with littermate controls (KO, N = 21; WT, N = 17), Distortion product otoacoustic emissions (DPOAE) thresholds at P30 are also significantly elevated, (KO, N = 8; WT, N = 6). ABR and DPOAE thresholds were increased at all tested frequencies (8, 11, 16, 22, and 32 kHz; Mann–Whitney U test, *p* < 0.001). **(F)** Scanning electron microscopy (SEM) images of WT and *Tmem30b* KO organs of Corti at P20. OHCs are pseudo-colored orange and IHCs yellow. Scale bars: 5 μm **(G)** High-magnification SEM images at P20 reveal stereocilia misorientation and OHC loss in *Tmem30b* KO mice, features absent in WT. Arrows point out areas of stereocilia misorientation. IHCs remain morphologically intact. Scale bars: 1 μm (left), 0.5 μm (middle and right). **(H)** Immunofluorescence images of organs of Corti from *Tmem30b* KO / ATP8B1-HA mice at P17 showing aberrant localization of ATP8B1-HA to the hair cell soma in the absence of TMEM30B (magenta). Scale bars: 5 μm

We next performed SEM analysis to assess the morphological defects in *Tmem30b* KO OHC hair bundles (**Fig. 5F**). Prior to overt cell loss, *Tmem30b* KO OHCs exhibited fractured and disorganized hair bundles. We did not observe the same hair bundle fusion that occurred in the *Atp8b1* KO OHCs, possibly pointing to a difference in cell deterioration between the two mouse lines (**Fig. 5G**). IHCs displayed no detectable morphological abnormalities. Overall, the degenerative phenotype in *Tmem30b* KO mice largely parallels that of *Atp8b1* KO mice, supporting a shared requirement for the ATP8B1-TMEM30B flippase complex in maintaining OHC integrity.

To determine how the loss of TMEM30B affects the localization of ATP8B1-HA, we examined ATP8B1-HA localization on the *Tmem30b* KO mouse background. In control mice, ATP8B1-HA is enriched in stereocilia, and not found below the hair bundle and the apical membrane. In contrast, in the absence of TMEM30B, ATP8B1-HA showed markedly increased somatic staining with reduced confinement to the hair bundle, indicating protein mislocalization (**Fig. 5H**). These findings demonstrate that TMEM30B is necessary for correct targeting of ATP8B1, providing a mechanistic explanation for the closely overlapping phenotypes observed in the two KO models.

### ATP8B1 and TMEM30B deficiency causes aberrant PS exposure in OHC bundles

ATP8B1 deficiency is predicted to impair PS internalization from the outer plasma membrane leaflet (Klomp et al. 2004). Persistent PS exposure could disrupt MET function and/or initiate downstream pathways that contribute to hair-cell degeneration as observed in both KO mouse models. We therefore examined whether *Atp8b1* and *Tmem30b* KO OHCs exhibit abnormal PS exposure. We performed Annexin V (AnV) labeling on apical turns of P15 organs of Corti (Zhang et al. 1997), a developmental timepoint that precedes overt hair cell loss in these mutants (Koopman et al. 1994) (**Fig. 6A, C**). AnV labeling was restricted to stereocilia bundle and apical membrane and not detected along the lateral membrane or within the cell body. Both *Atp8b1* KO and *Tmem30b* KO mice exhibited significantly elevated AnV fluorescence compared with wild-type controls (**Fig. 6B, D**). We occasionally observed weak AnV labeling also in IHCs, at the apical membrane (often appearing to be enriched at cell junctions) and the hair bundle (**Fig. 6A, C**). Together, these findings indicate that loss of ATP8B1-TMEM30B flippase activity leads to aberrant PS exposure predominantly in OHC stereocilia, preceding and likely contributing to hair cell degeneration in both KO models.

**Figure 6.**
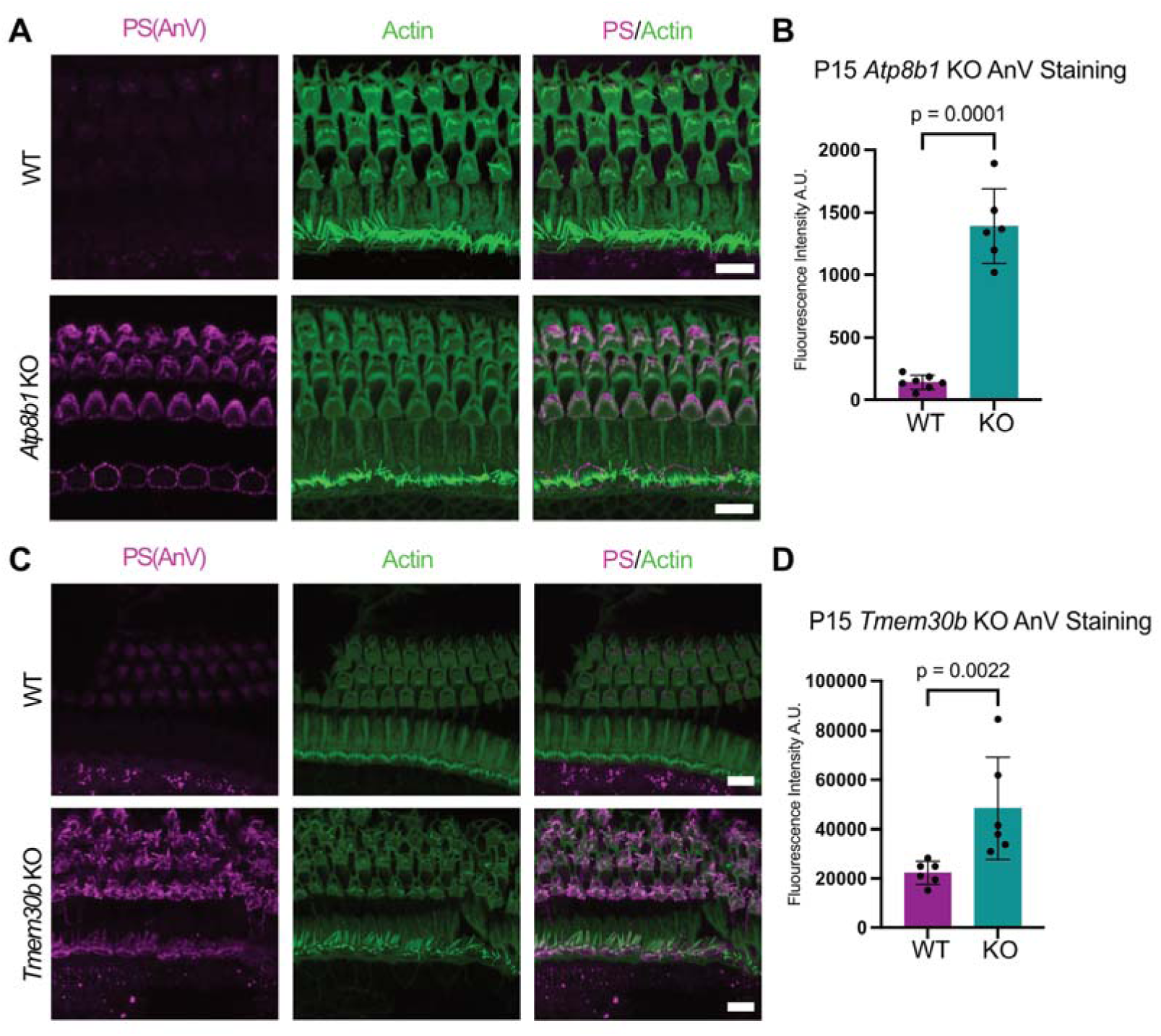
Loss of plasma membrane asymmetry precedes hair cell loss in *Atp8b1* and *Tmem30b* knockout mice. **(A)** Representative confocal images of WT and *Atp8b1* KO cochleae at P15 stained with Annexin V to label externalized phosphatidylserine (PS) and phalloidin to visualize F-actin. Increased Annexin V labeling is observed in *Atp8b1* KO hair cells prior to overt cell loss. Scale bar, 10 μm. **(B)** Quantification of Annexin V fluorescence intensity in WT and *Atp8b1* KO cochleae (n = 5–7 cochleae per group). *Atp8b1* KO hair cells show significantly increased PS externalization compared with WT (Welch’s *t*-test, *p* < 0.0001). **(C)** Representative confocal images of WT and *Tmem30b* KO cochleae at P15 stained with Annexin V and phalloidin. Increased Annexin V labeling is similarly observed in *Tmem30b* KO hair cells. In *Atp8b1* and *Tmem30b* KO organs, weak AnV labeling was occasionally observed also in IHCs, at the apical cell junctions and the hair bundle. Scale bar, 10 μm. **(D)** Quantification of Annexin V fluorescence intensity in WT and *Tmem30b* KO cochleae (n = 5–7 cochleae per group), demonstrating significantly elevated PS externalization in KO hair cells (Welch’s *t*-test, *p* < 0.01).

## Discussion

In this study, we identify P4-ATPase ATP8B1 and its accessory subunit TMEM30B as essential regulators of OHC survival and auditory function. Our findings establish a direct mechanistic link between OHC stereocilia membrane lipid asymmetry and hearing by demonstrating that disruption of a stereocilia-localized flippase complex leads to aberrant PS exposure, early hair-bundle pathology, progressive OHC degeneration, and rapid hearing loss. Together, these data reveal a previously underappreciated requirement for active lipid homeostasis in OHCs and provide cellular and mechanistic validation of TMEM30B as a novel deafness gene.

### A flippase complex is selectively enriched in OHC stereocilia

A central finding of this work is the highly selective localization of ATP8B1 and TMEM30B to OHC stereocilia. Among the P4-ATPases expressed in hair cells, ATP8B1 uniquely localizes to the hair bundle, was not detectably enriched in IHCs, and becomes progressively enriched in hair bundles during postnatal maturation, coinciding with the onset and maturation of hair cell MET. This specificity likely serves to accommodate the uniquely high mechanical and metabolic demands placed on OHC stereocilia. ATP8B1 is strongly enriched within stereocilia membranes, with minimal localization in the plasma membrane below the epithelial barrier level. Loss of TMEM30B results in mislocalization of ATP8B1, demonstrating that this β-subunit is required not only for flippase activity but also for proper targeting and retention of ATP8B1 in the stereocilia membrane.

### Lipid asymmetry as a potential structural and functional determinant of MET

Recent cryo-EM studies, together with the discovery that TMC1 and TMC2 possess intrinsic lipid-scrambling activity, have prompted a re-evaluation of MET in which membrane lipids and their asymmetric distribution are re-considered as active determinants of channel function rather than passive structural components (Ballesteros and Swartz 2022; Jeong et al. 2022; Clark et al. 2024; Peineau et al. 2025). Loss of the asymmetric distribution of membrane with their bulky and charged headgroups such as PS is known to have profound biophysical consequences for the cellular membrane (Kakuda et al. 2022; X. Li et al. 2024; Wang et al. 2025), and the stereocilia membrane will not be an exception. Redistribution of PS to the outer leaflet alters bilayer thickness (Wang et al. 2025), curvature stress, and membrane viscosity (Caputo et al. 2025), parameters that are particularly critical in stereocilia (George et al. 2020; George et al. 2023; George and Ricci 2025), where MET channel gating is exquisitely sensitive to membrane mechanics.

Building on this framework, a recent model proposed by George and Ricci suggests that TMC1/2 scramblase activity, by lowering membrane viscosity, reduces the energetic cost of channel gating, which may improve both the speed and sensitivity of the mechanotransduction response (George and Ricci 2025). If lipid scrambling is an integral feature of MET, however, a compensatory mechanism must exist to prevent unchecked collapse of membrane asymmetry. It is tempting to speculate that the ATP8B1-TMEM30B flippase complex fulfills this role by continuously restoring lipid asymmetry and maintaining OHC membranes within a “Goldilocks zone” of viscosity optimal for OHC MET.

This model is supported by the spatial distribution of ATP8B1, which is most abundant at the base of stereocilia and within the apical OHC membrane, with decreasing levels toward the stereocilia tips where MET occurs. Such a gradient suggests that TMC1-mediated lipid scrambling predominates at the site of MET, while ATP8B1-TMEM30B activity at the bundle base and apical membrane prevents widespread PS exposure, which could be interpreted as an “eat-me” signal and trigger pathological and phagocytic responses by surrounding supporting cells. Consistent with this model, elevated PS exposure in both *Atp8b1* and *Tmem30b* KO hair cells precedes overt structural degeneration, indicating that continuous flippase activity is required to preserve stereociliary membrane composition and stability.

### Selective requirement for stereocilia flippase activity in OHCs versus IHCs

A striking aspect of our finding is that in the cochlea, ATP8B1 is selectively expressed in OHC stereocilia and was not detected in IHCs under the conditions examined. One possible explanation is that IHCs stereocilia express another flippase. This seems unlikely as we systematically assessed all P4-ATPases expressed in hair cells and did not find evidence for an alternative stereocilia-enriched P4-ATPase in IHC bundles. Instead, we think it likely that IHCs, at least in the stereocilia, lack flippase activity altogether. It is possible that IHCs maintain lipid asymmetry using a different strategy, for example, by relying on a distinct set of lipid transporters localized outside the bundle, possibly sufficient given the lower MET resting open probability in IHCs (Johnson et al. 2011; Peng et al. 2013; Corns et al. 2014), resulting in lower basal scrambling pressure. It is also possible that beyond P4-ATPase flippases, complementary regulation could also arise from other enzymes controlling lipid abundance (synthesis and degradation) and compartmentalization (Blom et al. 2011).

### Early bundle pathology precedes hair-cell loss

Annexin V mediated detection of PS and structural analyses reveal that PS externalization and hair-bundle degeneration precedes hair-cell death in both KO models. Notably, IHCs remain morphologically and functionally intact, consistent with their lack of ATP8B1 and TMEM30B expression. This selective vulnerability underscores the unique lipid homeostasis requirements of OHC stereocilia, likely reflecting an adaptation to their higher MET activity and mechanical load. A recent study reported Annexin V labeling of hair bundles beginning around P7 in wildtype mouse OHCs, suggesting PS exposure on the outer leaflet during normal development (George and Ricci 2025). This observation contrasts with our findings, in which robust and consistent Annexin V labeling was observed only in *Atp8b1* and *Tmem30b* KO hair cells. We note, however, that we occasionally detected low-level Annexin V signal in control samples, even in IHCs (as shown in **Fig. 6**). Given the sensitivity of the live Annexin V labeling procedure to rapid deterioration of cochlear hair cells, particularly OHCs after ototoxic insult (Shi et al. 2007; Goodyear et al. 2008), this signal was initially interpreted as background or false-positive staining. Nonetheless, an alternative explanation is that OHC stereocilia possess higher PS exposure under normal physiological conditions, an expected phenomenon if TMC1/2 mediated MET causes phospholipid scrambling. Such low-level exposure of PS may fall near the detection threshold of Annexin V staining, especially if quickly reversed by flippase activity, and thus be difficult to resolve reproducibly. Further work will be required to determine whether regulated, developmentally controlled phospholipid exposure occurs in normal hair bundles and how it is balanced by flippase activity.

### Why do ATP8B1/TMEM30B-deficient OHCs die?

One notable feature of *Atp8b1* and *Tmem30b* KO phenotypes is their fast-progressing, base-to-apex pattern of OHC loss, reminiscent of several models in which MET is disrupted (e.g., TMC1/2 or CIB2 loss and related MET-deficient conditions) (Kawashima et al. 2011; Riazuddin et al. 2012; Zhao et al. 2014; Pan et al. 2018; Beurg et al. 2025). Loss of lipid asymmetry could directly compromise MET by altering bundle membrane mechanics and/or the availability of lipids on the inner leaflet that are required for normal MET. In this model, OHC degeneration would arise downstream of impaired MET signaling. At the same time, the OHC-specific degeneration observed in *Atp8b1* and *Tmem30b* KO mice also parallels, both in kinetics and cellular selectivity, the phenotype reported in mice with impaired stereocilia calcium extrusion via PMCA2 (Street et al. 1998; Kozel et al. 1998; Beurg et al. 2025). Loss of lipid asymmetry and/or altered phosphoinositide organization could reduce PMCA2 efficiency, compromising calcium clearance from stereocilia and promoting cytotoxic calcium accumulation. Consistent with this general possibility, PMCA activity is known to depend on membrane lipid composition. Specifically, PS on the inner leaflet was shown to be required for optimal PMCA activity (Zhang et al. 2009).

### Limitations of this study

A limitation of the present study is that loss of lipid asymmetry in OHC stereocilia was assessed primarily through PS externalization, as detected by Annexin V binding. Annexin V is a widely used indicator of disrupted membrane asymmetry, and the absence of PS exposure in WT mice, together with its robust appearance in ATP8B1- and TMEM30B-deficient OHCs, supports a role for ATP8B1 in maintaining membrane lipid organization. However, this approach specifically reports PS exposure and does not directly inform on the asymmetric distribution of other lipid species. ATP8B1 has been reported to transport PS in several experimental contexts, and our data are therefore fully consistent with the possibility that PS is a direct substrate of ATP8B1 in OHC stereocilia membranes. At the same time, independent evidence from cell-based, biochemical, and structural studies indicates that ATP8B1 can also transport phosphatidylcholine (PC) and phosphoinositides, with recent work demonstrating ATP8B1-mediated flipping of PI(4,5)P_2_ (Takatsu et al. 2014; Gómez-Mellado et al. 2022; Dieudonné et al. 2022; Bhandari et al. 2025). Consequently, the PS externalization observed in *Atp8b1* and *Tmem30b* KO mice may reflect loss of asymmetry of PS itself, of other ATP8B1 substrates, or of multiple lipid species simultaneously. Because robust in situ reporters for leaflet-specific PC or phosphoinositide distribution are currently limited, our study cannot directly distinguish among these possibilities. Thus, while PS exposure provides a clear and biologically meaningful readout of altered lipid asymmetry, it should not be interpreted as indicating that ATP8B1 function in stereocilia is exclusively or primarily PS-centric. Instead, our findings support a broader role for ATP8B1 in maintaining the asymmetric organization of stereocilia membranes, with PS externalization serving as a sensitive marker of this process. Future studies employing lipid-specific biosensors, reconstitution-based approaches, or advanced lipidomic strategies may be important for defining the full spectrum of ATP8B1 substrates in stereocilia and for determining how perturbations in distinct lipid species contribute to OHC structure and function.

### TMEM30B as a deafness gene and therapeutic implications

While ATP8B1 has been previously implicated in hearing loss, evidence was limited and mechanistic insight lacking (Stapelbroek et al. 2009). Our data provide strong validation of ATP8B1’s role in hearing and identify TMEM30B as an essential partner whose loss phenocopies ATP8B1 deficiency at molecular, cellular, and functional levels. These findings align with a previous large-scale mouse mutagenesis screen reporting hearing loss in *Tmem30b* KO mice (Bowl et al. 2017). Importantly, ATP8B1 and TMEM30B present attractive therapeutic features. Their expression begins postnatally and peaks after hearing onset in mice, and hair-cell loss in KO mice occurs after ∼P16, providing a potential window for intervention. Furthermore, TMEM30B is a single-exon gene of modest size (∼3.5 kb including UTRs), making it well suited for AAV mediated gene delivery and faithful recapitulation of endogenous expression.

### Conclusions and future directions

In summary, this study establishes ATP8B1 and TMEM30B as important regulators of membrane lipid asymmetry in OHC stereocilia and hair-cell survival, and points to a potentially essential role for lipid flipping in hair cell MET. Future studies will be required to quantify, using cellular electrophysiology, how lipid asymmetry influences MET channel gating and sensitivity, and to determine which additional lipid species are regulated by ATP8B1 in stereocilia.

## Materials and Methods

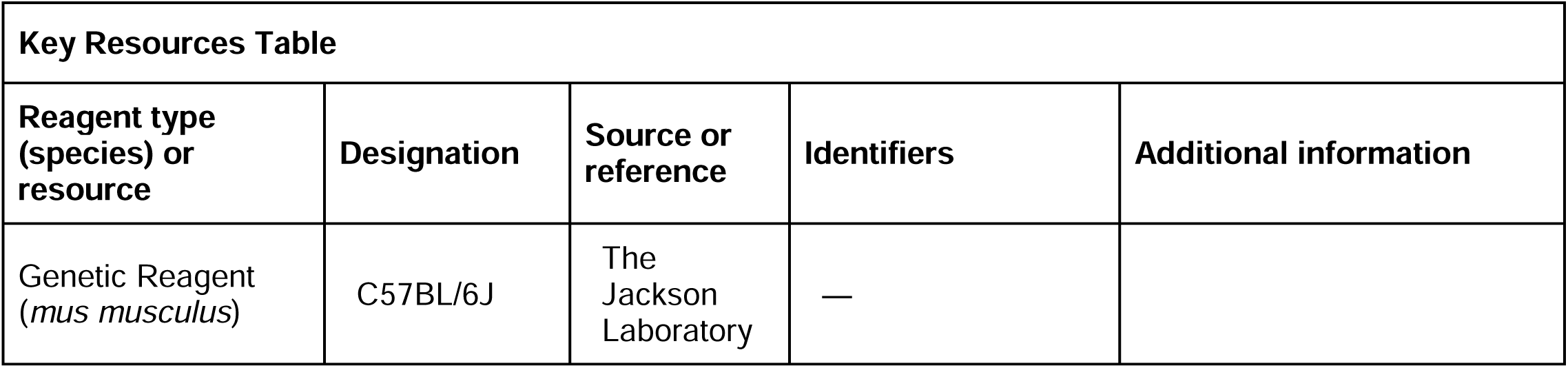
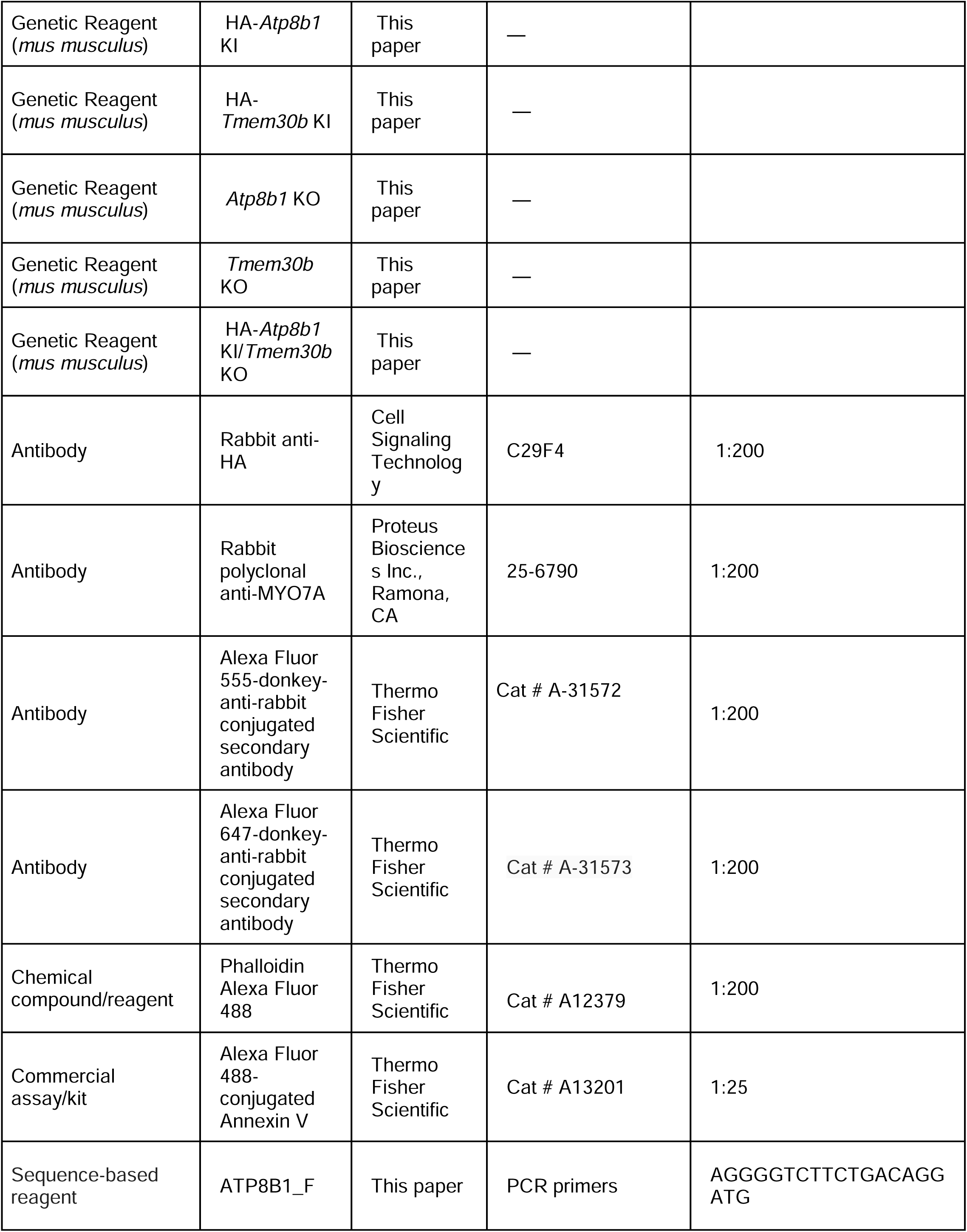
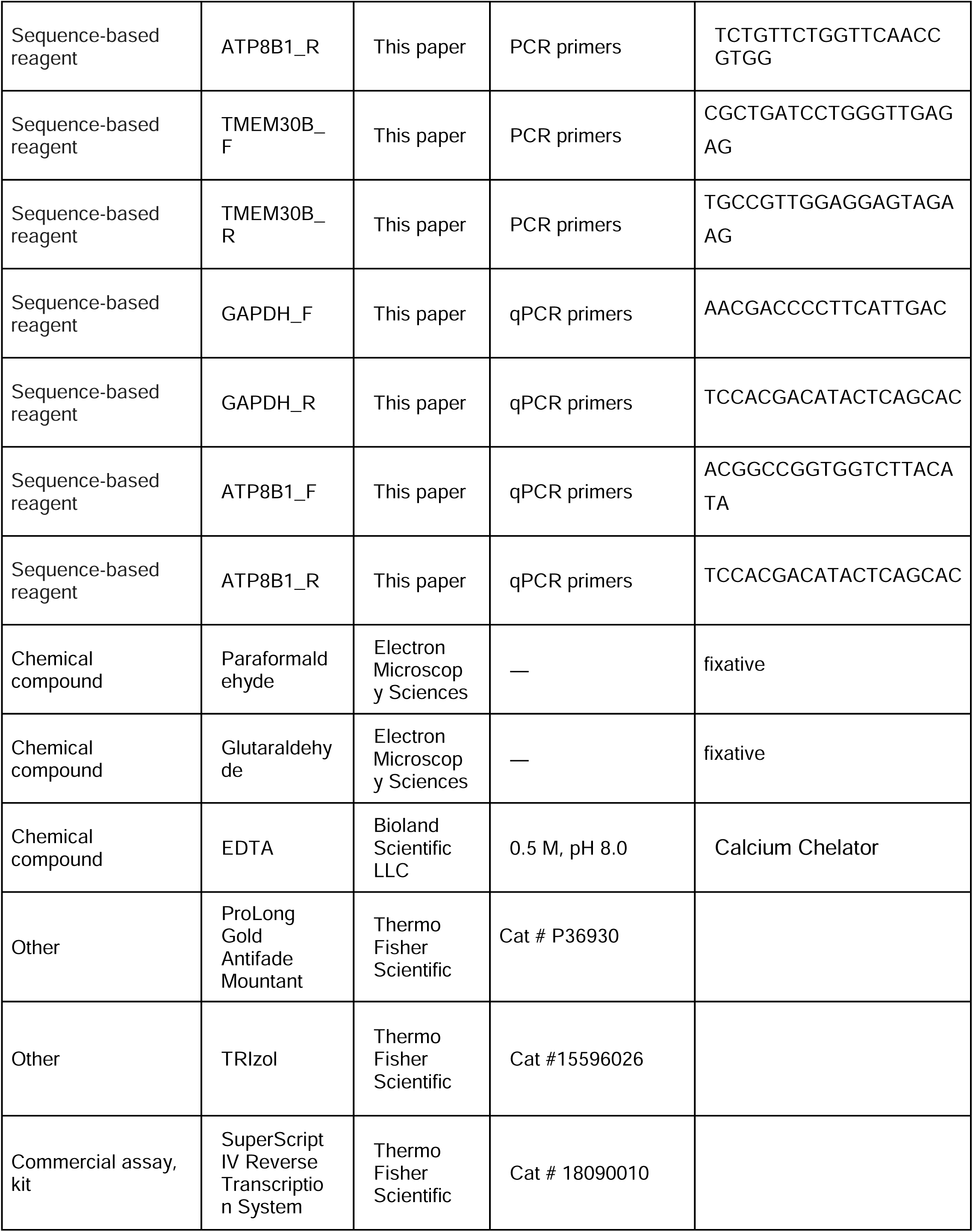
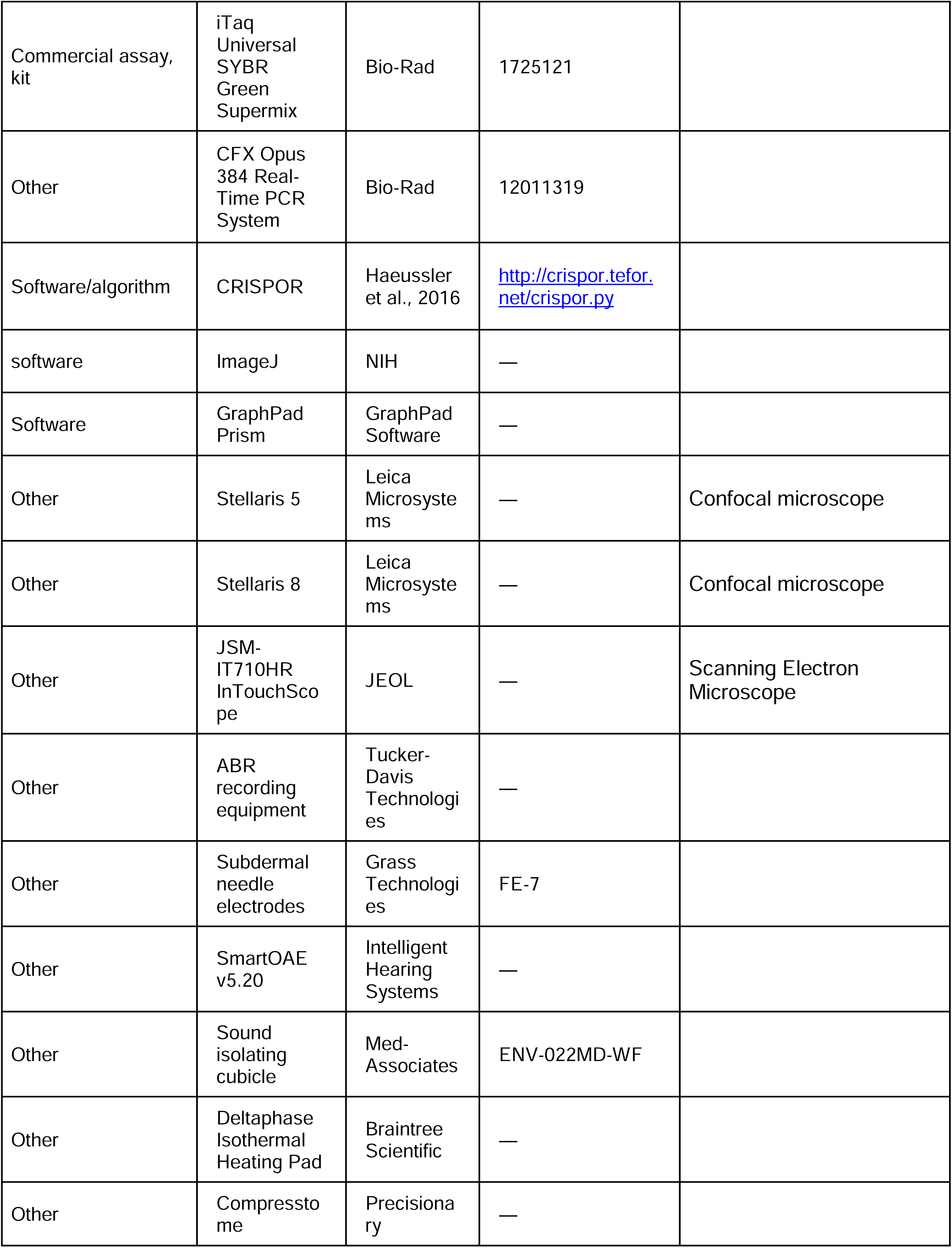
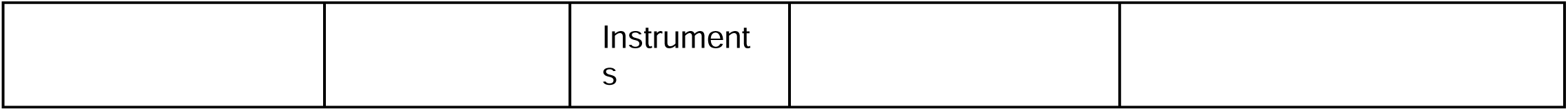

### Animal Care and Handling

All murine experiments conducted for this study adhered to the guidelines established by the University of Virginia Institutional Animal Care and Use Committee (IACUC) and the NIH. Animal protocols were reviewed and approved by the University of Virginia IACUC. Mice were housed in an AAALAC-accredited vivarium located in the basement of our research building. Animals were maintained on a 12-hour light/dark cycle, with cage changes performed every five days. Mice had ad libitum access to standard rodent chow and water provided through lixit systems. Control mice were C57BL/6J mice obtained from Jackson Laboratory. Experimental mice ranged from postnatal day 0 (P0) to P5 and were euthanized via rapid decapitation. Mice aged P6 and older were euthanized using CO2 asphyxiation, followed by cervical dislocation as a secondary method to ensure humane euthanasia.

### Generation of Experimental Mice

Genetically modified mouse models used in this study were generated using CRISPR/Cas9-mediated genome editing to create knockout or knock-in alleles of ATP8B1 and TMEM30B. Guide RNA target sequences were). CRISPR/Cas9 target sequences were designed using the online tool CRISPOR (http://crispor.tefor.net/crispor.py) to generate KI and KO alleles for *Atp8b1* and *Tmem30b*(Haeussler et al. 2016). The selected guide RNA target sequences were **AAGCACAGAAAGAGACT** for *Atp8b1* and **CGCCGTGGCGCTCCAGGTCA** for *Tmem30b*.

For *Atp8b1*, the genomic sequence surrounding the target site and illustrating the KO modification is shown below:

TCAGCCCAACACAGGAAAGTCACGACTCACTCCTGTTTCCTCCTTTTCTCTCCAGGTAGTTCCAATTTGCC AGCAGA**AT<u>G</u>**<u>AGCACAGAAAGAGACTCG</u>GAAACAACATTTGATGAGGAGTCTCAGCCCAACGATGAAGTG GTCCCCTACAGTGATGATGA

The HA epitope-coding sequence (TACCCATACGATGTTCCAGATTACGCT) was inserted immediately downstream of the start codon (ATG, bold). The KO allele was generated by deletion of 19 nucleotides (underlined).

For *Tmem30b,* the following CRISPR target sequence was identified: **CGCCGTGGCGCTCCAGGTCA**. The genomic sequence surrounding the target site and illustrating the KO modification is shown below:

GCGAGCGGAGGCGGGAGGCTCGCCGCTGATCCTGGGTTGAGAGTCGGCGGC<u>CGCAGGCCCCGGCTGA</u> <u>GATCCCCGCC**ATG**ACCTGGAGCGCCACGGCGCGGGGCG</u>CGCACCAGCCCGACAACACCGCCTTCACTC AGCAGCGCCTCCCCGCCTGGCAGCC

As for ATP8B1, the HA epitope-coding sequence was inserted immediately downstream of the start codon (ATG, bold). The KO allele was generated by deletion of 54 nucleotides (underlined).

### Genotyping

Genotyping of *Atp8b1* alleles yielded PCR products of 239 bp (wt), 266 bp (HA KI), and 220 bp (KO). Primers used were:

Forward: AGGGGTCTTCTGACAGGATG

Reverse: TCTGTTCTGGTTCAACCGTGG

Genotyping of *Tmem30b* alleles yielded PCR products of 232 bp (wt), 259 bp (HA KI), and 178 bp (KO). Primers used were:

Forward: CGCTGATCCTGGGTTGAGAG

Reverse: TGCCGTTGGAGGAGTAGAAG

### Mouse colony maintenance

*Atp8b1* KO mice were maintained as heterozygous breeders. All other lines (*Atp8b1* HA KI, *Tmem30b* KO, *Tmem30b* HA KI, HA-*Atp8b1* KI/*Tmem30b* KO) were maintained as homozygous breeding colonies.

### Immunofluorescence

Mice were sacrificed via decapitation (for P5 and younger) or CO2 asphyxiation followed by cervical dislocation (for P6 and older). Cochleae were carefully dissected and placed into a dish containing phosphate-buffered saline (PBS; GIBCO, Thermo Fisher Scientific). Under a dissecting microscope, the ossicles were removed, and the cochleae were transferred to a well containing 4% paraformaldehyde (PFA; Electron Microscopy Services). The cochleae were flushed through the oval and round windows using a transfer pipette to ensure proper fixation and incubated in the fixative for 20 minutes at room temperature. After fixation, the cochleae were washed three times for five minutes each in PBS. For P7 and younger mice, cochleae were immediately dissected. For P8 and older mice, cochleae were decalcified in 0.5 M ethylenediaminetetraacetic acid (EDTA; Bioland Scientific LLC), pH 8.0, for up to two weeks, depending on bone density. Organs of Corti were either dissected as whole mounts under a stereomicroscope using forceps and micro-dissection scissors or sectioned into 50 μm slices using a Compresstome (Precisionary Instruments, Ashland, MA). Dissected or sliced organs of Corti were transferred to blocking buffer and incubated for two hours at room temperature. The blocking buffer consisted of PBS supplemented with 1% bovine serum albumin (BSA), 3% normal donkey serum, and 0.2% saponin. Following blocking, tissues were incubated overnight at 4°C with primary antibodies diluted in the same blocking buffer. The following primary antibody was used: rabbit anti-HA (C29F4, Cell Signaling Technology, Danvers, MA).The next day, tissues were washed three times for five minutes each in PBS, then incubated with fluorescently labeled secondary antibodies and phalloidin in blocking buffer for two hours at room temperature (Alexa Fluor 555-conjugated secondary antibody, Thermo Fisher Scientific, Secondary-Cat# A-31572, Cat# A-31573, Phalloidin-Cat A12379. After secondary antibody incubation, tissues were washed three additional times in PBS, mounted onto glass slides using ProLong Gold Antifade Mounting Media (Thermo Fisher Scientific, P36930), and allowed to cure overnight. Confocal imaging was performed using a Leica Stellaris 5 confocal microscope equipped with a 63x oil-immersion objective.

### ATP8B1-HA Fluorescence Line Scan Analysis

Cochleae from ATP8B1-HA knock-in mice were immunostained with anti-HA antibody and labeled with phalloidin to visualize F-actin as described in the immunofluorescence protocol above. Confocal images were acquired using a Leica Stellaris 5 confocal microscope equipped with a 63× oil-immersion objective. High-resolution images of outer hair cell stereocilia were collected using identical acquisition parameters across samples. To assess sub-stereociliary localization of ATP8B1-HA, fluorescence intensity line scans were performed along individual stereocilia using ImageJ (NIH). Line scans were drawn manually along the length of both long (row 1) and short (row 2) stereocilia from the base at the cuticular plate to the stereocilia tip. Fluorescence intensity profiles for ATP8B1-HA and phalloidin were extracted and background-subtracted. For each stereocilium, positional distance was normalized from 0 (base) to 1 (tip), and fluorescence intensity was normalized to the total signal per stereocilium to allow comparison across cells. Normalized intensity profiles were averaged across stereocilia, and shaded regions represent the standard error of the mean (SEM).

### Fluorescence Intensity Quantifications of ATP8B1-HA and TMEM30B-HA

Cochleae were immunostained with anti-HA antibody and labeled with phalloidin as described in the immunofluorescence protocol above. Confocal images were acquired using a Leica Stellaris 5 confocal microscope equipped with a 63× oil-immersion objective. For each experiment, all images were acquired using identical acquisition parameters, including zoom and laser power, across genotypes within that experiment. Fluorescence intensity of the HA signal was quantified in ImageJ (NIH). Whole-mount images were resliced to generate orthogonal views, and regions of interest (ROIs) were manually drawn around the hair bundle and hair cell soma. Background fluorescence was measured in adjacent regions lacking specific signal and subtracted from all measurements. For all analyses, measurements were restricted to the middle cochlear turn. For each cochlea, fluorescence intensity was measured in five outer hair cells and averaged to generate a single biological replicate (n = 4–6 cochleae per genotype, 1 cochlea representing 1 mouse). Data analysis and statistical testing were performed using GraphPad Prism.

### Quantitative PCR

qPCR primers were designed to detect ATP8B1 expression levels. Total RNA was extracted using TRIzol reagent (15596026, ThermoFisher Scientific). Reverse transcription was performed using the SuperScript™IV Reverse Transcription system (18090010, ThermoFisher Scientific). qPCR reactions were conducted using iTaq Universal SYBR Green Supermix (1725121, Bio-Rad, CA, USA) on a CFX Opus 384 Real-Time PCR System (12011319, Bio-Rad) for fluorescence measurement. Transcript level of *Atp8b1* is normalized to the transcript level of *Gapdh*. Expression analysis was performed in GraphPad Prism.

*Atp8b1* Primers used were:

Forward: ACGGCCGGTGGTCTTACATA

Reverse: GCTTGTCACTCACGTCCTGAT

*Gapdh* Primers used were:

Forward: AACGACCCCTTCATTGAC

Reverse: TCCACGACATACTCAGCAC

### Hair Cell Counts

Cochleae were immunostained for MYO7A ( Rabbit polyclonal anti-MYO7A, Proteus Biosciences Inc., Ramona, CA, 25-6790) and labeled with phalloidin as described in the immunofluorescence protocol above. Confocal images were acquired using a Leica Stellaris 5 confocal microscope equipped with a 63× oil-immersion objective. For each experiment, all images were acquired using identical acquisition parameters, including zoom and laser power, across genotypes within that experiment. Hair cells were quantified using the Cell Counter plugin in ImageJ (NIH) based on the presence of MYO7A-positive cells. Counts were performed separately for apical, middle, and basal cochlear turns, defined by their relative position along the cochlear spiral (n = 3–7 cochleae per genotype, 1 cochlea representing 1 mouse). Data analysis and statistical testing were performed using GraphPad Prism.

### Annexin V Staining and Quantification

P15 mice were euthanized and cochleae were rapidly dissected and gently opened to facilitate Annexin V penetration. Cochleae were flushed with Alexa Fluor 488–conjugated Annexin V (Thermo Fisher Scientific Cat # A13201) diluted 1:25 (10 μL Annexin V in 250 μL L-15 medium, GIBCO, Thermo Fisher Scientific; total volume ∼260 μL) and then incubated for 30 min at room temperature. Following incubation, cochleae were washed twice in L-15 medium, fixed in 4% paraformaldehyde, and washed three times in PBS (5 min each). Samples were then dissected and mounted using ProLong Gold antifade mounting medium. Confocal images were acquired using a Leica Stellaris 5 confocal microscope equipped with a 63× oil-immersion objective for the *Atp8b1* knockout experiments and a Leica Stellaris 8 confocal microscope for the *Tmem30b* knockout experiments due to instrument availability. For each experiment, all images were acquired using identical acquisition parameters, including zoom and laser power, across genotypes within that experiment.Fluorescence intensity of the Annexin V signal was quantified in ImageJ. ROIs were manually drawn around individual outer hair cells, and mean fluorescence intensity was measured using the ImageJ (NIH) measurement function. Background fluorescence was measured in adjacent regions lacking specific signal and subtracted from all measurements. Analyses were restricted to the apical cochlear turn, as decalcification was not performed prior to Annexin V incubation. For each cochlea, fluorescence intensity was measured in five outer hair cells and averaged to generate a single biological replicate (n = 4–7 cochleae per genotype, 1 cochlea representing 1 mouse). Data analysis and statistical testing were performed using GraphPad Prism.

### Scanning Electron Microscopy

Mice were sacrificed via decapitation (P5 and younger) or CO2 asphyxiation followed by cervical dislocation (P6 and older). Cochleae were carefully dissected and placed in a dish containing phosphate-buffered saline (PBS; GIBCO, Thermo Fisher Scientific). Under a dissecting microscope, ossicles were removed, and the cochleae were transferred to conical tubes containing 2.5% glutaraldehyde and 2% paraformaldehyde (PFA; Electron Microscopy Services). Samples were incubated in this fixative solution overnight at 4°C. For adult mice, cochleae were subsequently incubated in 0.5 M ethylenediaminetetraacetic acid (EDTA; Bioland Scientific LLC), pH 8.0, for up to two weeks to facilitate decalcification, depending on bone density. Following decalcification, the cochleae were further dissected to expose the organ of Corti, ensuring the complete removal of the tectorial membrane. Samples were prepared for scanning electron microscopy using the OTOTO procedure (Davies and Forge 1987). This process included sequential osmium tetroxide and thiocarbohydrazide treatments, followed by dehydration through a graded ethanol series and critical point drying using hexamethyldisilazane (HMDS; Electron Microscopy Services). Once dried, samples were sputter-coated with a thin layer of platinum to enhance conductivity. Imaging was performed on a JEOL JSM-IT710HR InTouchScope™ Scanning Electron Microscope housed in our microscopy core facility. SEM images were pseudocolored using Affinity (Canva Pty Ltd.)

### Hearing Tests in Mice

Prior to Auditory Brainstem Response (ABR) tests mice were anesthetized with an intraperitoneal injection of 20 mg/kg ketamine hydrochloride (Covetrus Inc.) and 10 mg/kg xylazine hydrochloride (Covetrus Inc.). ABR tests were performed in a sound isolating cubicle (Med-Associates, product number: ENV-022MD-WF). To maintain body temperature mice are kept on a Deltaphase isothermal heating pad (Braintree Scientific). ABR recording equipment was purchased from Tucker Davis Technologies (Alachua, FL). Recordings were captured by subdermal needle electrodes (FE-7; Grass Technologies). The noninverting electrode was placed at the vertex of the midline, the inverting electrode over the mastoid of the right ear, and the ground electrode on the upper thigh. Stimulus tones (pure tones) were presented at a rate of 21.1/s through a high-frequency transducer (Tucker Davis Technologies). Responses were filtered at 300–3000 Hz and threshold levels were determined from 1024 stimulus presentations at 8, 11.3, 16, 22.4, and 32 kHz. Stimulus intensity was decreased in 5–10 dB steps until a response waveform could no longer be identified. Stimulus intensity was then increased in 5 dB steps until a waveform could again be identified. If a waveform could not be identified at the maximum output of the transducer, a value of 5 dB was added to the maximum output as the threshold. DPOAEs of the same group of WT and *Atp8b1* and *Tmem30b* KO mice were recorded. While under anesthesia for ABR testing, DPOAE was recorded using SmartOAE ver. 5.20 (Intelligent Hearing Systems). A range of pure tones from 8 to 32 kHz (16 sweeps) was used to obtain the DPOAE for the right ear. DPOAE recordings were made for f2 frequencies from 8.8 to 35.3 kHz using a paradigm set as follows: L1 = 65 dB, L2 = 55 dB SPL, and f2/f1 = 1.22. Mice were separated by sex when tested for initial ABR and DPOAE experiments. We did not find any significant difference in ABR and DPOAE thresholds based on sex (not shown). ABR analysis for ATP8B1-HA, TMEM30B-HA*, Atp8b1* Ko, and *Tmem30b* Ko mice contain both males and females. DPOAE analysis for *Atp8b1* Ko, and *Tmem30b* Ko mice contain both males and females.

### Statistics and Reproducibility

All statistical analyses were performed using GraphPad Prism (GraphPad Software, La Jolla, CA, USA). Auditory brainstem response (ABR) thresholds were compared between genotypes using Mann–Whitney U tests. For all statistical analyses, significance was defined as *p* < 0.05. In figures, statistically significant *p*-values are indicated, whereas non-significant comparisons are not shown. Unless otherwise indicated, error bars represent standard deviation. Analyses in this study use biological replicates. Experiments were repeated between two and four times. Power analyses were conducted using preliminary means and standard deviations to determine appropriate sample sizes. Based on an effect size of Cohen’s *d* = 1.4 calculated from preliminary ABR data, a one-tailed test with α = 0.05 and a desired power of 0.8 (80%) indicated that a sample size of 10 mice per group (20 total) was sufficient to detect statistically significant differences between genotypes (Faul et al. 2007, 2009). Sample sizes for immunofluorescence experiments (hair cell counts, HA fluorescence intensity, and Annexin V staining) were determined based on prior experience with similar experiments, as the variability between genotypes could not be estimated a priori.

## Data and Resource Availability

Raw data used to create this manuscript are available on https://doi.org/10.17605/OSF.IO/GQBMF. Mouse lines and other reagents are available from the corresponding author upon request.

## Acknowledgments

We thank Wenhao Xu and Daniel Grigsby at Genetically Engineered Murine Model Core (University of Virginia, RRID:SCR_025473) for generating mouse models used in this study. We thank Sijie Hao from Advanced Microscopy Facility (University of Virginia, RRID: SCR_018736) for the support of SEM imaging. Graphics were created in Biorender. Funding sources: J.B.S is supported by NIH grant RO1DC018842 and R01DC021176-03. H.N.D is supported by NIH grant F30DC023139-01.

